# Open-Rosalind: Tool-First Biomedical LLM Agents with Process-Aware Benchmarking

**DOI:** 10.64898/2026.05.06.722404

**Authors:** Liang Wang

## Abstract

Large language models are increasingly used as scientific agents, yet the flexibility that benefits general-purpose agents can conflict with the accountability required in biomedical research. We study whether biomedical agents can be organized around auditable constraints rather than unconstrained autonomy. We present **Open-Rosalind**, a tool-first bio-agent system designed around four operational principles: evidence-grounded outputs, trace completeness, workflow-constrained execution, and explicit tool mediation for factual claims. To evaluate these principles, we introduce **Open-Rosalind BioBench**, a process-aware benchmark that measures not only task accuracy but also tool correctness, citation presence, trace completeness, and failure rate.

On a strict in-house benchmark, the reference pipeline achieves 81.4% accuracy with complete execution traces. In multi-model ablations and paired replications, removing tools reduces accuracy by 19.3 to 26.4 percentage points, indicating that tool-first execution is the strongest and most stable contributor to performance. Constrained workflows also reduce lower-tail failures for models that are weak at free-form tool use.

However, an author-independent 30-task hold-out initially revealed severe external-validity collapse on the deployment model. After diagnosing five routing and normalization failures and applying targeted fixes, hold-out accuracy improved from 17.8% to 53.3%, and the most concerning negative comparison against a `no_tool` baseline disappeared. These results position Open-Rosalind as a biomedical-agent study with an explicit external-validity audit, rather than as a claim that protocol constraints alone guarantee superior performance.

## 1 Introduction

Within a span of two years, large language models have evolved from text completion engines into autonomous agents that plan, invoke external tools, and iterate over multi-step tasks. In domains such as software engineering and general information work, this transition has produced clear gains, in part because mistakes are rapidly visible and easily corrected. Biomedical and life-science research impose a different standard. The problem is not capability: modern models can read a UniProt entry, parse a FASTA sequence, and reason about substitutions. The problem is *epistemic accountability*.

A scientific claim is not merely an assertion; it is a binding to a verifiable source. When a general-purpose LLM agent confidently reports that a protein is located on a particular chromosome arm, or cites a PubMed identifier in support of a mutation pathogenicity claim, the reader has no operational way to distinguish a tool-grounded fact from a hallucinated one. Worse, the agent may itself fail to draw this distinction, freely paraphrasing tool outputs alongside parametric knowledge in a single fluent paragraph. For wet-lab researchers, clinicians, or anyone whose downstream decisions are guided by such answers, this opacity is a fundamental defect, not an implementation gap.

The position taken in this paper is that bio-agent systems require a different kind of design contract than general-purpose agents:

*In biomedical AI, the question is not whether an agent can produce an answer, but whether the answer can be traced, audited, and replayed*.

Pursuing this contract leads to four design constraints that shape every layer of the system: factual content must come from explicit tools rather than parametric memory; every claim in an answer must be cited to a tool-derived source; every step of execution must be recorded in a replayable trace; and the agent must operate within bounded, pre-declared workflows rather than free-form planning loops. Each of these constraints sacrifices some flexibility in exchange for an auditable invariant.

This paper studies **Open-Rosalind**, a tool-first bio-agent system built around these constraints, together with a prototype system, a benchmark suite, and an external-validity repair study. We frame the work as three interlocking components. The first is a *principled architecture* (Sections 3–4) that codifies the four design principles into a layered stack in which atomic tools, MCP-style tool interfaces, deterministic skills, a hybrid router, an agent, and a bounded multi-step harness each carry their own reproducibility invariant. The second is a *prototype system* (Section 5) that demonstrates the architecture end-to-end on real biological tasks: sequence analysis, protein annotation, literature retrieval, and mutation assessment. The third is **Open-Rosalind BioBench** together with controlled ablations and a hold-out audit (Sections 6–7), designed not merely to maximize raw task accuracy but to measure whether bio-agent systems satisfy the framework’s reproducibility guarantees and how those guarantees behave off-distribution.

### Scope of claims

We do not argue that protocol discipline alone makes biomedical agents universally better, nor do we present Open-Rosalind as a finished product paper. The paper’s empirical claim is narrower. First, on the in-house benchmark, the tool-first design is a measurable accuracy lever relative to `no_tool`. Second, constraint appears to reduce lower-tail failures for models that are weak at unconstrained tool orchestration. Third, an external hold-out reveals that these gains are conditional on routing robustness and tool coverage: before repairs, the system collapses badly; after repairs, the most concerning negative comparison disappears, but a gap to ReAct remains. We therefore present the work as a biomedical-agent study with an honest diagnosis-and-repair cycle, rather than as proof that constrained bio-agents dominate more flexible alternatives in every setting.

## 2 From General Agents to Bio-Agents

The literature on LLM agents has converged on a small set of recurring patterns: *ReAct*-style interleaved reasoning and acting, *tool-use* protocols, *planner-executor* decompositions, and recursive task decomposition (e.g., AutoGPT-style loops). These patterns share a common philosophical commitment: *flexibility is a virtue*. The agent decides which tool to invoke, when to stop, when to backtrack, and how to summarize. This commitment is well-suited to open-ended environments, where the cost of failure is low and the diversity of tasks resists pre-specification.

Biomedical research is not such an environment. The set of legitimate workflows is small and well-defined: annotate a protein, search the literature on a topic, evaluate the impact of a mutation, compare two sequences. The cost of an unverifiable answer is high. The expectation that another researcher can reproduce the analysis is not optional. Under these conditions, flexibility is not a virtue but a liability: every additional degree of freedom afforded to the agent expands the surface over which errors and ambiguities can enter the result. Figure 1 illustrates the key architectural differences between general-purpose agents and reproducible bio-agents.

**Figure 1.**
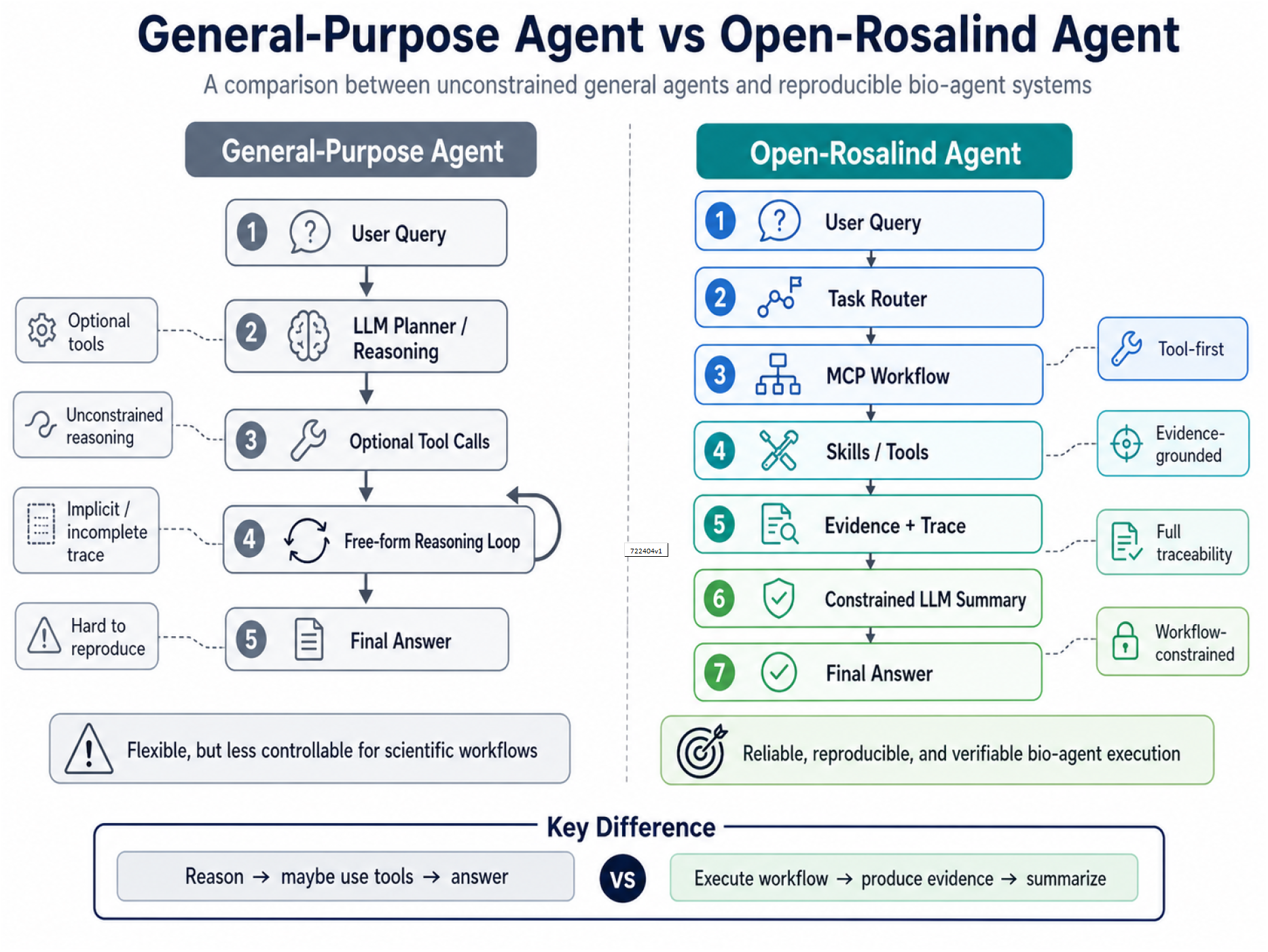
Architectural comparison: general-purpose agents vs. reproducible bio-agents. General agents prioritize flexibility with free-form reasoning and optional tool use; bio-agents prioritize accountability with mandatory tool-first execution, evidence grounding, and complete traceability.

Table 1 contrasts the design assumptions of general agents with those required for bio-agents. The right column of Table 1 should not be read as a list of capabilities omitted from general agents, but as a set of *guarantees added*. A bio-agent gains less behavioral freedom than a general agent but offers stronger contracts: every fact has a source, every step is logged, every workflow terminates within a known bound. The remainder of this paper develops these guarantees into a concrete framework.

**Table 1:** Design contrasts: general-purpose agents vs. reproducible bio-agents.

| Dimension | General-purpose agent | Bio-agent (this work) |
| --- | --- | --- |
| Knowledge source | LLM parametric memory + optional tools | Tools only; LLM is a synthesizer |
| Planning | Free-form, recursive, open-vocabulary | Templated, bounded, closed-vocabulary |
| Tool invocation | Optional, opportunistic | Mandatory for all factual claims |
| Output format | Natural language, untyped | Typed evidence + cited summary |
| Failure mode | Best-effort answer, possible hallucination | Explicit “insufficient evidence” |
| Trace | Best-effort logs (if any) | Mandatory, replayable, structured |
| Step bound | None or LLM-decided | Hard cap, externally enforced |
| Evaluation | Task accuracy | Accuracy + tool correctness + evidence + trace + failure rate |

## 3 Design Principles

The framework rests on four principles, summarized in Figure 2. Each principle is operational — it is enforced by a specific component of the system — rather than aspirational.

**Figure 2.**
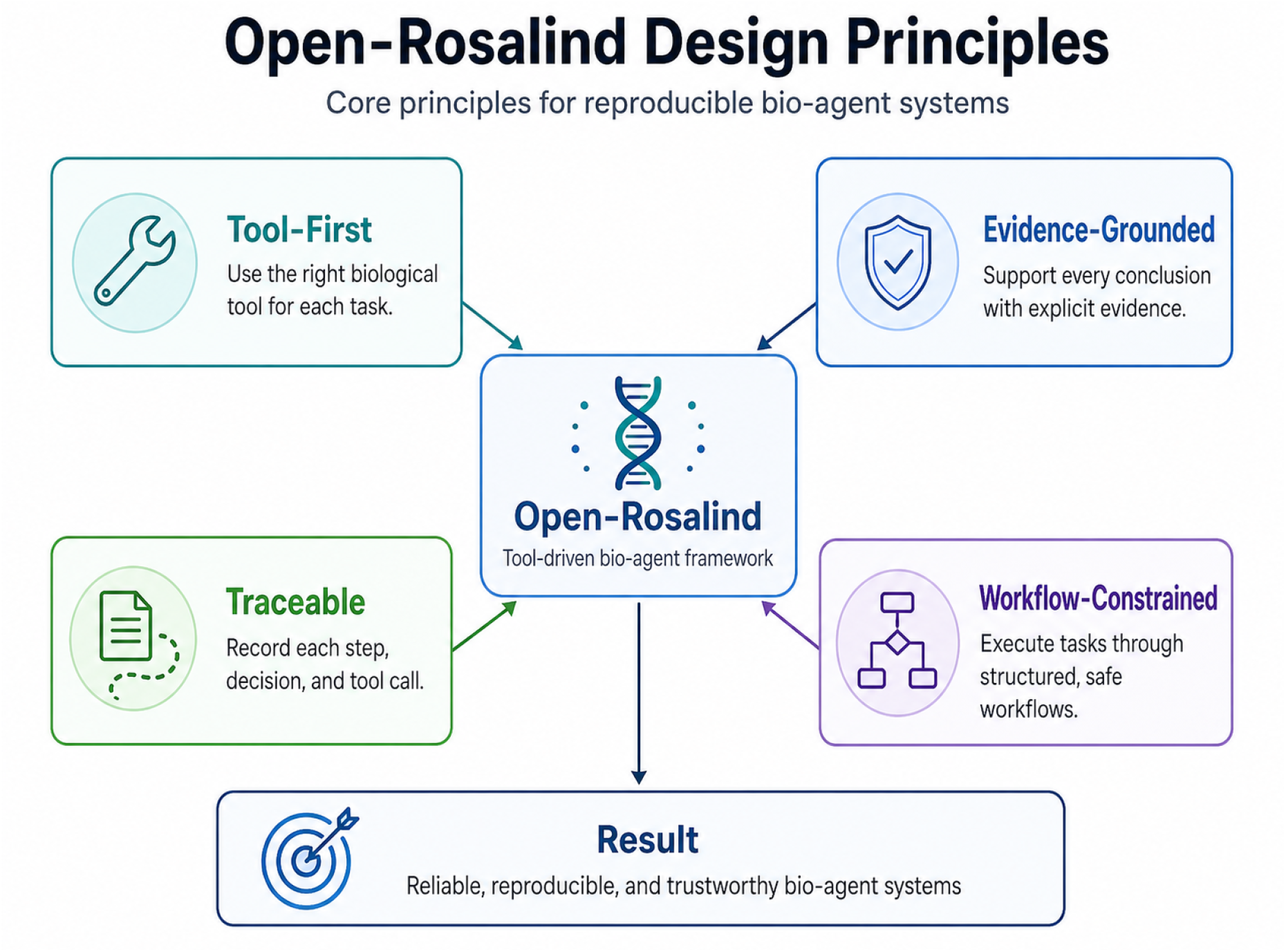
The four operational principles of Open-Rosalind. Each principle constrains the LLM’s role and is enforced by a specific layer of the architecture.

### 3.1 Tool-First Execution

The LLM never serves as a knowledge oracle. Every fact, computation, or annotation that appears in the final answer must originate from an explicit tool invocation registered in the system. The LLM is invoked only after tools have produced structured outputs, and its role is reduced to synthesizing those outputs into a natural-language summary. This re-positions the model as a *reader* rather than a *source*, and aligns the system with how scientific writing operates in practice: claims are made by citing data, not by recalling lore.

### 3.2 Evidence-Grounded Outputs

Every factual claim in the LLM’s output must carry an inline citation to a tool-derived source — a UniProt accession, a PubMed identifier, a structured statistic produced by an analysis routine. When the supporting evidence is insufficient, the LLM must say so explicitly rather than fall back on parametric guesses. This principle converts the conventional opacity of LLM outputs into something closer to a scientific paragraph: each sentence is either bound to a citation or marked as a deliberate non-answer.

### 3.3 Trace Completeness

Every routing decision, every tool invocation, every observed return value, and the final LLM prompt are recorded in a structured, replayable trace. The trace is not a debugging aid but a first-class artifact, intended to support reproduction by external readers and auditing by reviewers. The system treats a missing trace as a correctness failure on par with a wrong answer.

### 3.4 Workflow-Constrained Execution

Tasks are executed through pre-declared workflows rather than free-form agent loops. Single-step queries dispatch to one of a small fixed set of skills via a deterministic router; multi-step queries execute through one of a small fixed set of templated plans, under a hard step bound. The agent is not permitted to invent new tools, restructure the plan mid-run, or recurse beyond the bound. This sacrifices flexibility deliberately: predictability and termination are valued above adaptability.

## 4 Framework Architecture

Open-Rosalind organizes the four principles into a layered architecture in which each layer enforces a specific reproducibility invariant. We describe the layers from the bottom up.

### 4.1 Atomic Tools and MCP Interfaces

At the lowest level sit atomic tools — thin Python wrappers around external biological resources (UniProt, PubMed via Entrez) and local computations (BioPython sequence statistics, mutation diff). Atomic tools are intentionally minimal: each performs a single operation, returns a structured object, records latency and status, and never branches based on natural-language input. Tools are exposed to higher layers through MCP-style [1] interfaces, which provide a uniform contract for tool description, invocation, and result serialization. This boundary serves two purposes. First, it makes the system’s external dependencies explicit and version-controlled: a tool change is an interface change. Second, it allows alternative implementations (e.g., a local UniProt mirror, a cached PubMed index) to be substituted without changes to higher layers.

### 4.2 Skills as Composed, Deterministic Pipelines

A *skill* is the next layer up: a deterministic pipeline that composes one or more atomic tools to produce a typed output addressing a single biological question. The four skills used in this work — `sequence_basic_analysis, uniprot_lookup, literature_search, mutation_effect` — collectively cover the recurring questions in routine bioinformatics: *what is this sequence?, what is this protein?, what does the literature say?, what does this mutation do?*.

Skills are deliberately narrower than the LLM’s apparent reasoning capability. Their narrowness is the point: a skill is a unit of behavior that is small enough to test exhaustively, version explicitly, and reuse across both single-step and multi-step execution paths. Each skill carries a structured manifest declaring its category, safety level (safe / network / compute), the atomic tools it depends on, an input/output schema, and a small set of tested examples. Adding a new skill amounts to declaring its manifest and providing a handler; the rest of the framework discovers and integrates it without code changes.

### 4.3 Hybrid Router

The router maps an incoming user query to a skill (single-step path) or to a multi-step plan (harness path). It is intentionally hybrid: a rule-based pre-filter handles the unambiguous cases — a FASTA header, a UniProt accession pattern, a literature query containing the word “papers” — and an LLM-assisted intent classifier handles only the residual cases that mix natural language with embedded sequences or identifiers. The classifier is constrained to return a label from the small fixed skill set; it cannot invent new skills, and a parsing failure causes a deterministic fallback to the rule-based result.

This hybrid design reflects a recurring tension in agent systems: rule-based routing is brittle but auditable, while LLM-based routing is flexible but opaque. Open-Rosalind resolves the tension by giving rules first refusal: the LLM is consulted only when the rules cannot decide, and even then only inside a closed-vocabulary contract.

### 4.4 Single-Step Agent

The agent layer wraps the routed skill with the LLM-as-synthesizer step. Once the skill has executed and produced its typed output, the agent constructs an evidence-only prompt and asks the LLM to produce a Markdown summary with inline citations. The agent enforces the evidence-grounded-outputs principle through prompt design (the system prompt forbids claims without citations) and through post-hoc validation (claims without recognizable evidence markers trigger a fallback answer). The agent is also responsible for in-session context: a sliding window of prior turns is supplied to the LLM, allowing follow-up questions such as “what species is this protein in?” to be resolved against the prior annotation.

### 4.5 Multi-Step Harness

The harness orchestrates queries that require more than one skill. Rather than expose the agent to arbitrary planning, the harness chooses from a small fixed set of templates — in the present implementation: `protein_research` (sequence analysis → annotation → literature), `literature_review` (literature only), and `mutation_assessment` (mutation → annotation → literature). Each template is a sequence of skill invocations with explicit slots for entities (e.g., {protein_name}) that are filled from prior step outputs.

The harness enforces three invariants. First, the planner cannot invent new templates; it can only select from the registered set. Second, the runner observes a hard step bound (default five) that terminates execution regardless of the plan. Third, the per-step trace is structurally identical to a single-step trace, so multi-step workflows are reproducible and auditable in the same way as single-step queries.

The combined data flow — query, mode selection, single-step or harness path, skill execution, evidence and trace generation, and final answer — ensures that every execution path respects the framework’s four principles.

## 5 System Realization

We realized the proposed architecture as a compact experimental system consisting of a Python backend, a React-based front-end, and an embedded SQLite store for sessions, messages, and traces. The implementation is intentionally lightweight, because the scientific point is not scale but whether the protocol can be enforced end-to-end without a heavyweight orchestration stack. Figure 3 summarizes the resulting design.

**Figure 3.**
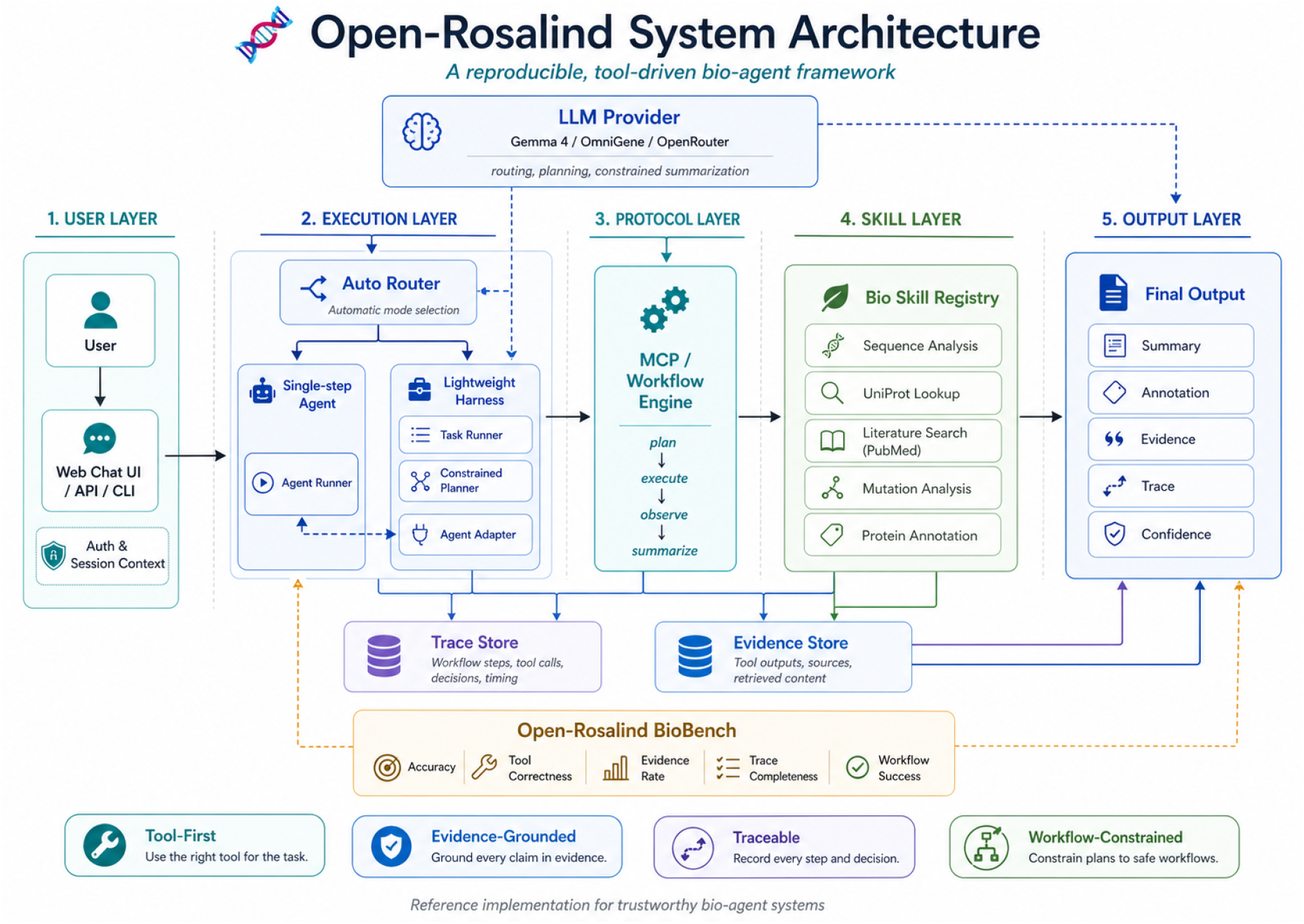
Open-Rosalind system architecture. The web UI communicates with a unified chat endpoint that dispatches to either single-step agent or multi-step harness paths. Both paths invoke skills from the registry, which in turn call atomic tools. All executions produce structured traces stored in SQLite.

**Figure 4.**
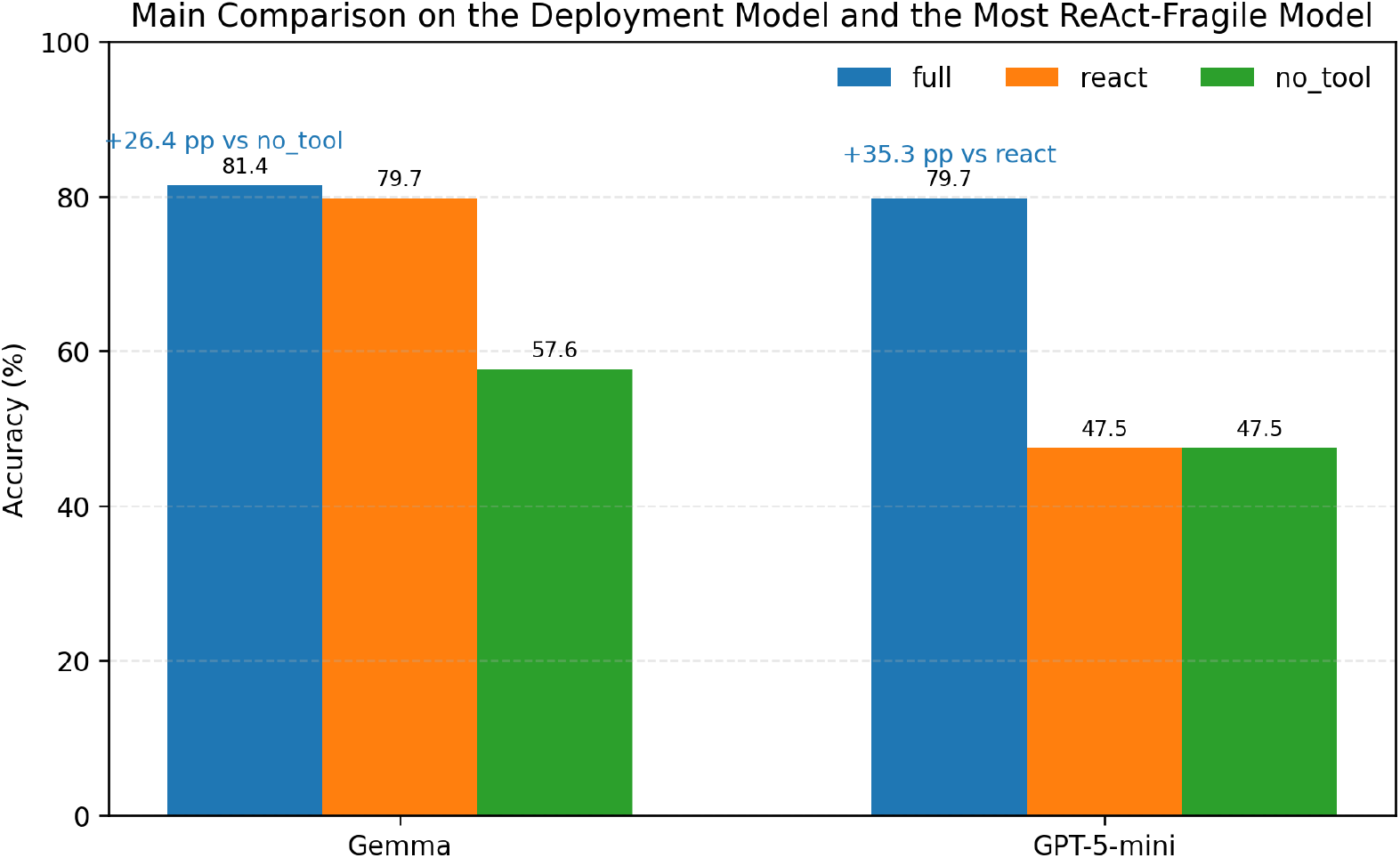
Main comparison emphasized in the text. The deployment target (Gemma) and the most ReAct-fragile model (GPT-5-mini) capture the paper’s two strongest claims: `full` substantially outperforms `no_tool` on both models, and it most clearly outperforms react when unconstrained tool use is unstable.

**Figure 5.**
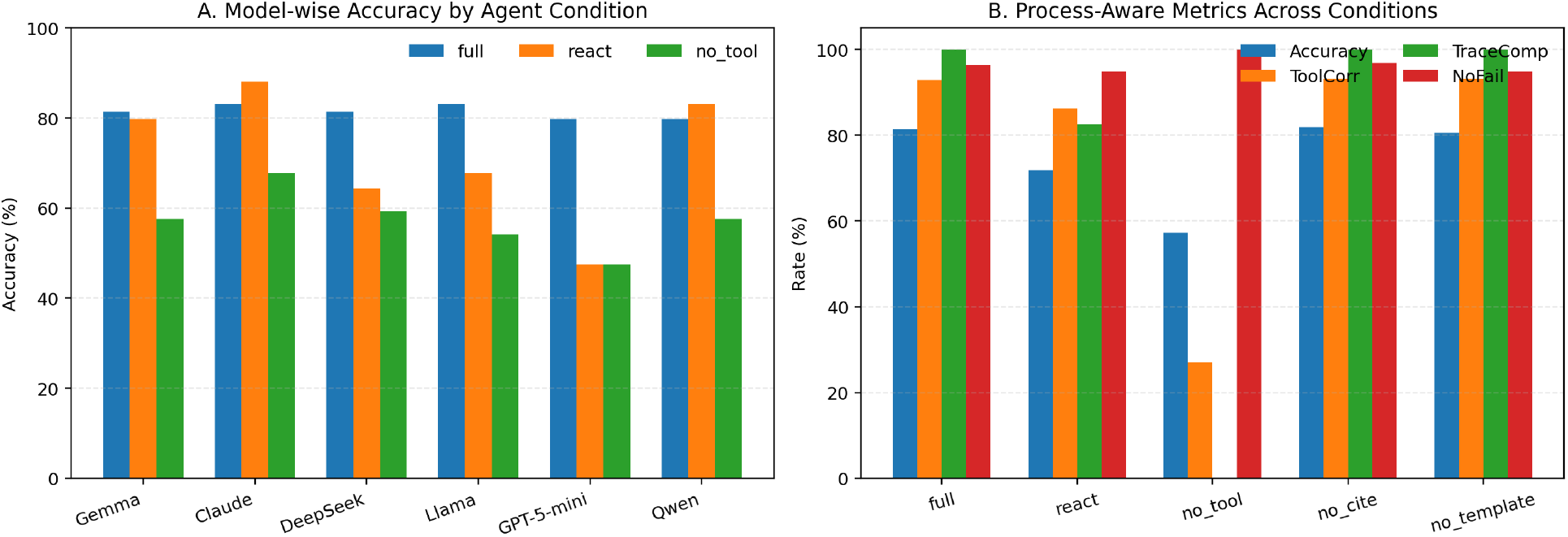
Experimental results beyond the architectural diagrams. Left: accuracy of `full`, `react`, and `no_tool`across six model families, showing that the constrained pipeline most clearly improves the lower-performing tool-use settings. Right: process-aware benchmark metrics aggregated across all models and conditions, illustrating that `no_tool` collapses tool correctness and trace completeness even when it remains failure-free.

**Figure 6.**
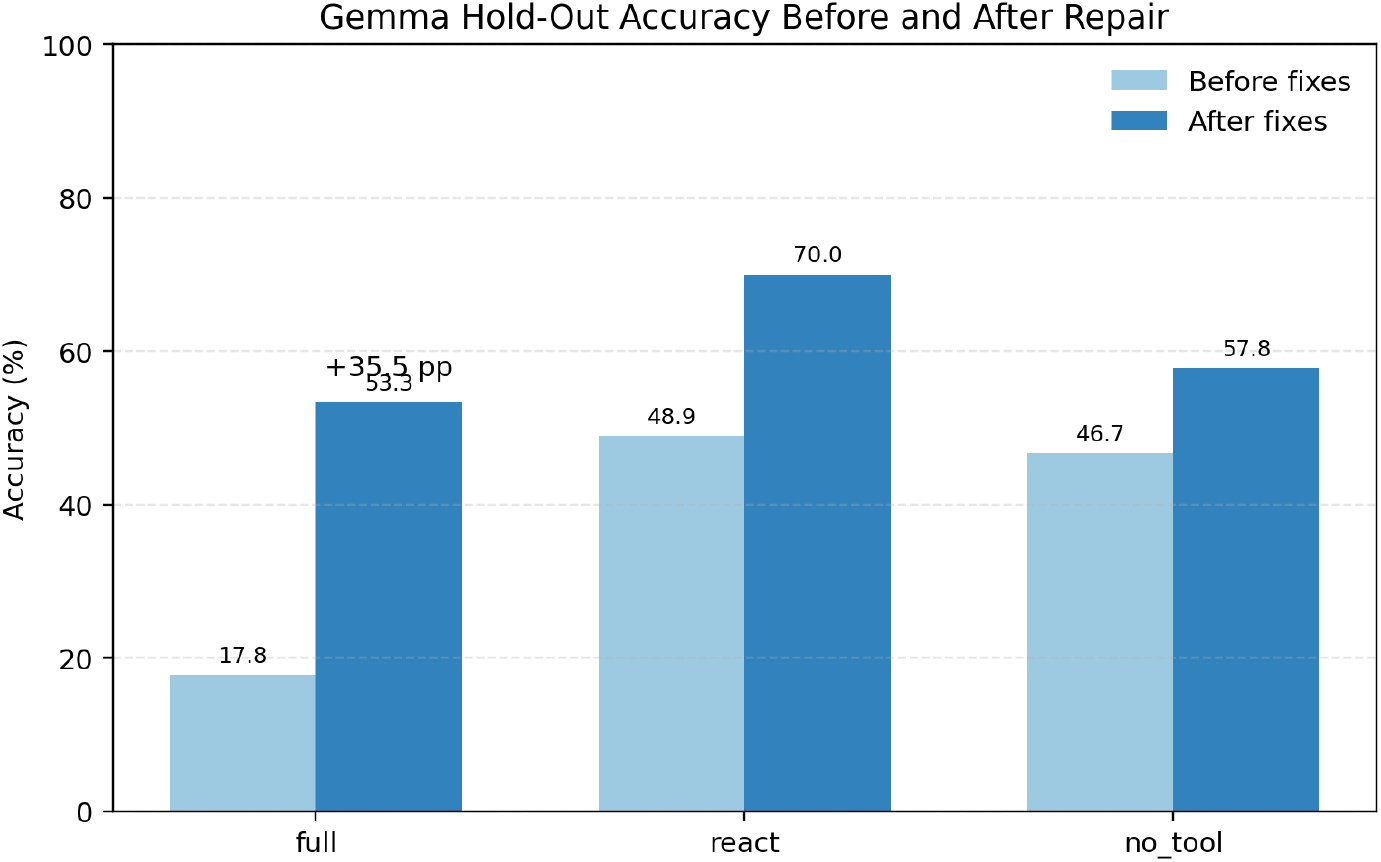
Gemma hold-out accuracy before and after the routing and normalization repairs. The most important change is not simply the gain in `full` accuracy, but the disappearance of the strongly negative `full` vs. `no_tool` comparison that had undermined the original claim.

The model layer is exchangeable: the system communicates with any OpenAI-compatible endpoint (OpenRouter, vLLM, Azure, or on-premise inference servers). The experiments reported here use `google/gemma-4-26b-a4b-it` via OpenRouter, but no part of the method is specific to that model. Skills are loaded from declarative manifests at startup, and the front-end presents a unified chat interface whose execution path is chosen by the router.

In addition to the chat interface, the system records skill registry state, session history, and per-session traces in forms that can be inspected after the fact. These records are essential to the trace-completeness principle: every interaction should leave behind material that can be reviewed, replayed, and audited.

## 6 Open-Rosalind BioBench

Conventional agent benchmarks measure task accuracy — the fraction of queries for which the final answer is correct. For auditable bio-agents, accuracy is necessary but not sufficient: an answer that happens to be right but is delivered through hallucinated tool calls, or with no recoverable trace, still fails the framework’s contract. **Open-Rosalind BioBench** is designed to make this distinction empirically tractable.

### 6.1 Splits and Provenance

The benchmark spans four task categories aligned with the four built-in skills: sequence analysis, protein annotation, literature retrieval, and mutation assessment. The repository currently contains 91 task instances across three splits: **Basic** (canonical inputs), **Edge** (mixed-modality or follow-up inputs), and **MultiStep** (queries requiring two or more skills). For the stricter model-comparison experiments in Section 7, follow-up variants are collapsed to 59 unique benchmark roots (32 Basic, 17 Edge, 10 MultiStep) so that each root question contributes once to paired statistical tests. Tasks are drawn from real biological objects — canonical UniProt accessions, PubMed-indexed publications, and proteins with documented variants — so that correctness can be checked against authoritative sources rather than model-generated ground truth.

### 6.2 Metric Design

Five metrics are reported per run. *Task accuracy* measures whether the final answer satisfies the task rubric. *Tool correctness* measures whether the realized skill or harness template matches the expected execution path. *Citation presence* measures whether factual claims carry recognizable evidence markers. *Trace completeness* measures whether the saved trace contains routing decisions, tool calls, statuses, and synthesis prompts. *Failure rate* measures crashes and timeouts separately from wrong answers. Because citation presence is partly syntactic, we additionally performed a manual validity audit on 20 sampled responses: 19/20 had all cited claims materially supported by the cited source, while 1/20 contained an over-broad paraphrase of a UniProt function field.

### 6.3 Why a Process-Aware Benchmark

The benchmark is intended to separate two questions that are often conflated in agent papers: whether a system returns the right answer, and whether it arrived there through a path that can be audited. The latter matters disproportionately in biomedical settings, where a wrong answer with a complete trace can be diagnosed and replayed, but a fluent answer with no trace cannot.

## 7 Empirical Results

### 7.1 Internal Benchmark Under a Strict Scorer

We begin with experiments using `google/gemma-4-26b-a4b-it`. Earlier development dashboards scored the 91 turn-level tasks with a looser keyword-overlap rule and reported 100.0% / 93.9% / 90.0% accuracy on Basic / Edge / MultiStep. For the paper’s main analyses, we instead use a stricter scorer on the 59 unique benchmark roots because it discriminates more sharply between conditions and supports paired statistical tests. The strict-scorer results are summarized in Table 2.

**Table 2:** Strict-scorer results on 59 unique BioBench roots using the reference Open-Rosalind pipeline on Gemma. Trace completeness remains saturated, but accuracy drops materially on Edge inputs, making the benchmark more discriminative than the original loose dashboard scorer.

| Split | Tasks | Accuracy | Tool corr. | CitePres | Trace / No-fail |
| --- | --- | --- | --- | --- | --- |
| Basic | 32 | 87.5% | 100.0% | 3.1% | 100.0% / 100.0% |
| Edge | 17 | 64.7% | 76.5% | 0.0% | 100.0% / 88.2% |
| MultiStep | 10 | 90.0% | 100.0% | 0.0% | 100.0% / 100.0% |
| **Combined** | **59** | **81.4%** | **93.2%** | **1.7%** | **100.0% / 96.6%** |

Two patterns matter. First, the Edge split is the real stress test: mixed natural language, embedded identifiers, and follow-up phrasing produce substantially lower accuracy than canonical inputs. Second, trace completeness remains saturated even when accuracy drops, meaning the framework localizes many failures to routing or factual synthesis rather than losing the execution record entirely. The weak point is citation presence under the strict per-sentence check: the current prompt tends to concentrate citations in a final evidence block rather than inline prose, so the metric functions here as an architectural target rather than a saturated success number.

### 7.2 Six-Model Ablations and Paired Replications

To test whether the framework’s behavior depends heavily on one underlying model, we ran a five-condition ablation across six LLM families: the `full` Open-Rosalind pipeline, a free-form ReAct baseline using the same atomic tools, a `no_tool` condition, a `no_cite` condition, and a `no_template` condition. The aggregate results over 1,770 runs are summarized in Table 3.

**Table 3:** Aggregate six-model ablation over 59 unique tasks and five conditions (1,770 runs total). The strongest and most stable effect is the drop from `full to no_tool`; citation and template ablations change accountability properties much more than average accuracy.

| Condition | Accuracy | Tool corr. | CitePres | TraceComp | NoFail |
| --- | --- | --- | --- | --- | --- |
| `full` | 81.4% | 92.9% | 0.3% | 100.0% | 96.3% |
| `react` | 71.8% | 86.2% | 0.6% | 82.5% | 94.9% |
| `no_tool` | 57.3% | 27.1% | 0.3% | 0.0% | 100.0% |
| `no_cite` | 81.9% | 93.2% | 0.0% | 100.0% | 96.9% |
| `no_template` | 80.5% | 93.2% | 0.3% | 100.0% | 94.9% |

The cleanest signal is the value of tools. Across models, removing tools collapses accuracy from roughly 80% to the mid-50s and drives tool correctness down to near-chance overlap with task keywords. To test whether this effect survives a corrected unit of analysis, we ran paired seed replications on the deployment target (Gemma) and on GPT-5-mini, which was the most ReAct-fragile model in the six-model sweep. Under cluster-aware permutation tests, the `full` pipeline beats `no_tool` by 26.4 percentage points on Gemma and 19.3 percentage points on GPT-5-mini (both *p* ≤ 10^−4^). The comparison to ReAct is more nuanced. On GPT-5-mini, the constrained pipeline beats ReAct by 35.3 percentage points and reduces catastrophic failures from 84.7% to 39.0%. On Gemma, however, `full` and ReAct are statistically indistinguishable on the in-house benchmark (*p* = 0.74). The support for the framework is therefore strongest as a *tool-first* and *lower-tail stability* claim, not as a universal claim that constrained workflows dominate free-form agents on raw accuracy.

### 7.3 External Hold-Out, Diagnosis, and Repair

To probe external validity beyond the in-house benchmark, we generated a 30-task hold-out set authored by an independent LLM and re-ran three paired conditions (`full, react, no_tool`) on Gemma and GPT-5-mini. Table 4 reports the Gemma results because Gemma is the deployment target and because the failure was most consequential there.

**Table 4:** Gemma results on the 30-task author-independent hold-out before and after the routing/normalization repairs. The most important change is that `full` is no longer significantly worse than `no_tool` after the fixes, although it remains below ReAct.

| Setting | full | react | no_tool |
| --- | --- | --- | --- |
| Before fixes | 17.8% | 48.9% | 46.7% |
| After fixes | 53.3% | 70.0% | 57.8% |

The initial hold-out materially changed the scientific interpretation of the system. On Gemma, the constrained pipeline collapsed from internal-benchmark performance to 17.8% accuracy and was significantly worse than both ReAct (Δ = −31.1 percentage points, *p* = 0.004) and `no_tool` (Δ = −28.9 percentage points, *p* = 0.002). Trace inspection showed that this was not a mysterious model failure. Most of the gap came from five concrete issues: stop-words such as “information” and “characterize” being preserved in search queries; natural-language mutation forms such as “KRAS G12D” failing to trigger the mutation workflow; Greek-vs-ASCII mismatches in keyword matching; embedded sequences going undetected when wrapped in prose; and insufficient coverage of literature keywords such as “publications” or “studies”.

After targeted fixes to the router, query cleaning, and scorer normalization, Gemma improved by 35.5 percentage points to 53.3%. More importantly, the most concerning negative comparison disappeared:`full` vs. `no_tool` became statistically indistinguishable (*p* = 0.72, 95% CI [−23.3, +13.3]). The gap to ReAct narrowed but remained significant (Δ = −16.7 percentage points, *p* = 0.032). This repair cycle clarifies the right interpretation of Open-Rosalind. Routing robustness is *necessary* for external validity, and several of the original failures were patchable engineering faults. The remaining gap, however, reflects a real design tradeoff: a tool-first system refuses to invent functional claims that its current tools cannot support. We therefore treat the hold-out not as an embarrassment to hide, but as evidence that bio-agent papers should report diagnosis-and-repair cycles rather than only polished in-distribution numbers.

## 8 Discussion

### Position relative to general-purpose agent frameworks

ReAct [20] and its descendants assume that flexibility is intrinsically valuable. For research domains in which the set of legitimate tasks is small, the cost of incorrect outputs is high, and reproducibility is non-negotiable, this assumption inverts. Open-Rosalind is in part an argument that the bio-agent literature should adopt a different default: that flexibility be unlocked deliberately, and that every degree of freedom granted to the agent be paid for in additional verification machinery.

### Position relative to biomedical LLMs

A complementary line of work — BioGPT [7], BioMedLM [6], Med-PaLM [5] — improves the LLM’s biomedical knowledge through specialized pre-training. Open-Rosalind is orthogonal to this line: the framework seeks to restrict what a model may assert without evidence. The experiments suggest that a disciplined protocol can raise the performance floor of smaller models, but they do not imply that protocol alone substitutes for model capability in every regime.

### Position relative to recent bio-agent systems

The past year has seen rapid progress in LLM-based biomedical agents. **Biomni** [8] is a general-purpose biomedical AI agent that unifies reasoning, retrieval, and code execution across diverse biological domains (genomics, immunology, microbiology, neuroscience), emphasizing breadth and autonomous task execution through free-form planning. **GeneAgent** [9] addresses hallucination in gene-set analysis by autonomously interacting with biological databases to verify its own output, introducing a self-verification loop that reduces parametric recall errors. **STELLA** [10] employs a multi-agent architecture with an evolving template library and dynamic tool ocean, enabling self-improvement through iterative refinement. A recent comprehensive review [11] surveys the landscape of LLM agents for biomedicine, highlighting methods, evaluations, and challenges across clinical and research applications.

Concurrent work has also addressed reproducibility and evaluation. **R-LAM** [12] proposes reproducibility-constrained large action models for scientific workflow automation, introducing structured action schemas and deterministic execution traces. Several benchmarks have emerged: an **AI Agent Evaluation Suite for Bioinformatics** [13] evaluates frontier models across multiple agent harnesses using LLM-based graders; a **Scientific Research Suite** [14] provides rigorous benchmarking for AI agents in scientific contexts; and frameworks for **multi-agent scientific AI evaluation** [15] propose process-level metrics, scaling curves, and adversarial robustness tests. A Nature Methods perspective [16] surveys the broader landscape, noting the transformative potential of autonomous systems while emphasizing the need for verification mechanisms.

Open-Rosalind’s distinct contribution relative to these systems is its focus on *protocol discipline as a first-class design constraint*. Where Biomni and STELLA prioritize capability breadth and self-evolution, and GeneAgent prioritizes self-verification, Open-Rosalind prioritizes *reproducibility guarantees enforced at every architectural layer*: every fact must originate from a tool, every step must be logged, every workflow must terminate within a known bound. The hold-out repair cycle in

Section 7.3 shows both the value and the limit of this position. These guarantees are not automatic — they require careful routing robustness and enough tool coverage to support the requested claims. We therefore view recent bio-agent systems as complementary: some explore what autonomous bio-agents *can do*; Open-Rosalind asks what they *must guarantee* to be trusted in scientific workflows.

### Limitations

The benchmark is still small and partly internal: the 91 task instances and 59 unique benchmark roots were authored during development, and the external hold-out, while independent of the system author, was still LLM-authored rather than human-authored. The harness templates are hand-coded; while their fixity is a feature for reproducibility, it bounds the range of workflows the system can express. The literature skill depends on PubMed availability and abstract coverage, and the mutation skill currently supports sequence-grounded assessment more strongly than mechanistic functional prediction. Finally, the hold-out results show that external validity remains only partially recovered: after repairs, the most worrying negative comparison disappears, but the full-vs-ReAct gap remains on Gemma.

### Future work

Three directions are immediate. First, expanding tool coverage for the failure classes exposed by the hold-out: richer mutation-effect predictors, broader literature retrieval, and ontology-backed protein annotation fields. Second, replacing the current internal benchmark with a human-authored blind split and multi-rater scoring protocol. Third, pushing reproducibility one step further by caching external tool responses and pinning model versions more tightly, so that traces become not only auditable but closer to bit-level replayable.

## 9 Conclusion

Open-Rosalind shifts the design of bio-agent systems from free-form reasoning to bounded, traceable workflow execution. Its main contribution is best read as a biomedical-agent study with an honest external-validity audit: a layered architecture with explicit invariants, a concrete experimental realization, a process-aware benchmark, and a diagnosis-and-repair cycle that makes the limits of the approach visible rather than implicit. The internal experiments support tool-first execution as a real accuracy lever, especially against `no_tool` baselines, and support constrained workflows as a lower-tail stabilizer for models that are weak at free-form tool use. The external hold-out, however, shows that these advantages are conditional on routing robustness and tool coverage. We hope the framework, the benchmark, and the released traces are useful not only as research artifacts, but as a concrete template for how biomedical agent papers can make their accountability claims measurable.

## Code, Data, and Reproducibility

The system, benchmark tasks, hold-out data, and specification are released open-source under the MIT license at https://github.com/maris205/open-rosalind. Per-task traces from the experiments reported here are included in the repository. The current release supports strong auditability through saved routing decisions, tool I/O, statuses, and synthesis prompts; exact replay still depends on live external APIs and hosted model versions, which we list as next-step work rather than a solved property.

